# The contrasting roles of nitric oxide drive microbial community organization as a function of oxygen presence

**DOI:** 10.1101/2021.12.09.472001

**Authors:** Steven A. Wilbert, Dianne K. Newman

## Abstract

Microbial assemblages are omnipresent in the biosphere, forming communities on the surfaces of roots, rocks, and within living tissues. These communities can exhibit strikingly beautiful compositional structures, with certain members reproducibly occupying particular spatiotemporal microniches. Despite this reproducibility, we lack the ability to explain these spatial patterns. We hypothesize that certain spatial patterns in microbial communities may be explained by the exchange of redox-active metabolites whose biological function is sensitive to microenvironmental gradients. To test this, we developed a simple community consisting of synthetic *Pseudomonas aeruginosa* strains with a partitioned denitrification pathway: a strict consumer and strict producer of nitric oxide (NO), a key pathway intermediate. Because NO can be both toxic or beneficial depending on the amount of oxygen present, this system provided an opportunity to investigate whether dynamic oxygen gradients can tune metabolic cross-feeding and fitness outcomes in a predictable fashion. Using a combination of genetic analysis, controlled growth environments and imaging, we show that oxygen availability dictates whether NO cross-feeding is deleterious or mutually beneficial, and that this organizing principal maps to the microscale. More generally, this work underscores the importance of considering the contrasting and microenvironmentally tuned roles redox-active metabolites can play in shaping microbial communities.

## Introduction

Given the diversity of microbial metabolic strategies and that microbes rarely live alone, the potential for metabolic interactions is vast ^1^. Metabolic interactions can be positive or negative and can have profound effects on the surrounding environment, from contributing to nutrient flux to stimulating the immune system of an infected animal or plant. Numerous ecological models have been developed to better understand how such interactions influence the growth of a community ^2–4^. However, these models typically focus on simple growth kinetics involving a shared or exchanged beneficial metabolite and do not consider how changing microenvironments over small spatial scales might influence the initiation and/or quality of the interaction. Yet it is a well-documented ecological phenomenon that microbes are confronted with environmental changes that can tune these interactions, such as shifting oxygen concentrations, in both space and time ^5–8^. How such spatiotemporal environmental changes impact the function of exchanged metabolites is often overlooked in these models.

Diffusible, redox-active metabolites, be they organic or inorganic, are particularly relevant to consider in this context, as these molecules’ reactivity is highly sensitive to local chemistry. Certain redox-active metabolites have the potential to be both toxic (i.e. generate reactive oxygen species under oxic conditions) or beneficial (i.e. contribute to energy conservation under anoxic conditions), thereby having the potential to play dynamic roles in the development of a microbial community ^9^. Such contrasting effects may be particularly important when the metabolite is an intermediate in a common metabolic pathway. For example, if two microbes are reliant on each other for growth due to an intermediate that can be toxic to the producer but is essential for the consumer, it stands to reason that their growth and subsequent environmental impact will depend on microenvironmental conditions that affect the properties of the exchanged intermediate. The potential for variable local interactions due to shifting environmental gradients would therefore be expected to determine when, where, and how certain microbial interactions would manifest spatiotemporally.

As a step toward testing this hypothesis, we sought to create a simple synthetic model system in which to explore the effect of the microenvironment on the interactions between two bacterial partners linked via an exchanged redox-active metabolite. Four criteria guided our development of such a system. First, we required that the metabolite be “agathokakological”, a word derived from Greek roots meaning “composed of good and evil”. In other words, a metabolite that is predictably contrasting: beneficial or harmful according to its environmental context. Second, we needed to be able to measure and perturb the most important environmental parameter—oxygen—that affects its physiological impacts. Third, we aimed to control the production and utilization of this metabolite via the generation of mutant strains from an isogenic background in order to focus strictly on the capacity to make or use the metabolite. Finally, we sought a metabolic exchange that is of broad ecological relevance across many environments.

The exchange of nitric oxide (NO) met all of our criteria. NO is known in both the eukaryotic and microbial worlds as a potent signaling molecule and a highly reactive nitrogen species that can damage a variety of cellular components ^10^. In the absence of oxygen, NO is toxic at high concentrations by antagonizing Fe-S centers and causing DNA damage; yet in the presence of oxygen, much lower levels of NO can be toxic due the generation of peroxynitrite ^11,12^. Microbes have numerous mechanisms to deal with low levels of NO toxicity such as flavohemoglobin proteins (Fhp) that can reduce NO to nitrous oxide (N_2_O) or mediate its oxidation to nitrate (NO_3_^-^) depending on the environment ^13,14^, and other dioxygenases can sequester NO ^15,16^. However, as its concentration rises, and if producing cells or neighbors are unable to reduce it to nitrous oxide (N_2_O), it reaches toxic levels and leads to local growth inhibition or death ^17,18^.

NO production is also a landmark step in denitrification, where the soluble substrate nitrite (NO_2_^-^) is reduced to gaseous NO. Denitrification *sensu stricto* plays a critical role in the loss of nitrogen in soils ^19–21^. If an organism contains a full pathway to sequentially reduce nitrate (NO_3_^-^) > NO_2_^-^ > NO > N_2_O > nitrogen gas (N_2_), harmless N_2_ will be released into the environment where it can eventually become recycled by nitrogen fixing species ^22^. However, the complete reduction of NO_3_^-^ to N_2_ may occur as a polymicrobial community effort in which different members contain only a subset of the denitrification pathway ^23–26^. Synthetic pseudomonads have previously been shown to exchange nitrite under strictly anoxic conditions, leading to predictable spatial patterns that are sensitive to pH-dependent abiotic nitrite reduction to toxic nitrogen species ^27–30^. In an independent study, environmental isolates that naturally differ in their possession of denitrification genes were used to create a model that predicted interactions via denitrification intermediates^23^; yet, in this case, NO production was only considered for its potential toxicity rather than any potential for energy conservation. Because PA uses a NO reductase that does not directly contribute to proton translocation (qNor), it is often considered only for its detoxifying potential ^10^. However, physiological studies have shown that NO reduction in other, qNor-containing strains can contribute to ATP synthesis as the sole terminal electron acceptor (TEA) ^31,32^. To our knowledge, the ability for NO to be exchanged between cells in a manner that supports growth has not been demonstrated.

Here, we utilized the denitrifying organism *P. aeruginosa* strain PA14 to test the hypothesis that the spatial interactions of strains exchanging NO can be predictably altered as a function of oxygen in the microenvironment. *P. aeruginosa* is a cosmopolitan organism, notorious for being an opportunistic human pathogen, but is also found in sediments and soils ^13^. *P. aeruginosa* is also a classic denitrifier that encodes all enzymes necessary for the complete reduction of nitrate to nitrogen gas and can be easily genetically manipulated, permitting the creation of a synthetic NO cross-feeding community. Our findings provide a conceptual lens through which to interpret more complex microbial community patterning in nature and disease involving the exchange of redox active metabolites.

## Results

### Design and characterization of a synthetic NO cross-feeding community

The PA14 genome contains a full suite of respiratory enzymes for the complete dissimilatory reduction of NO_3_^-^ to N_2_ (Nar, Nap, Nir, Nor, Nos) (Figure 1A) ^13^. Previous studies have shown that these steps occur sequentially and thus intermediates can build up and leak from the cell ^33^. Because the PA14 NO reductase (Nor) does not directly contribute to proton translocation and in fact requires the consumption of protons for its reaction, it is often dismissed for contributing to energy conservation. However, if we trace the flow of electrons (Figure 1B, gray dashed lines) originating from the oxidation of NADH by the Nuo NADH dehydrogenase, through the quinone pool to the bc_1_ complex, a total of six protons are translocated prior to NO reduction ^34^. If we account for the two protons consumed by the reduction of two NO molecules to one N_2_O and one H_2_O (Figure 1B, solid line), four protons still remain per NADH oxidized that could contribute to the proton motive force. Electron flow to the N_2_O reductase (Nos), which reduces N_2_O to N_2_, is also thought to follow the same pathway ^34^. Thus, when considering the full electron transport chain, we expect for each NADH oxidized, a net of 4H^+^ to be translocated per NO reduced all the way to N_2_. Accordingly, there is good reason to expect NO to be able to support sufficient energy conservation to power growth under anoxic conditions, provided its toxicity under these conditions is managed.

**Figure 1.**
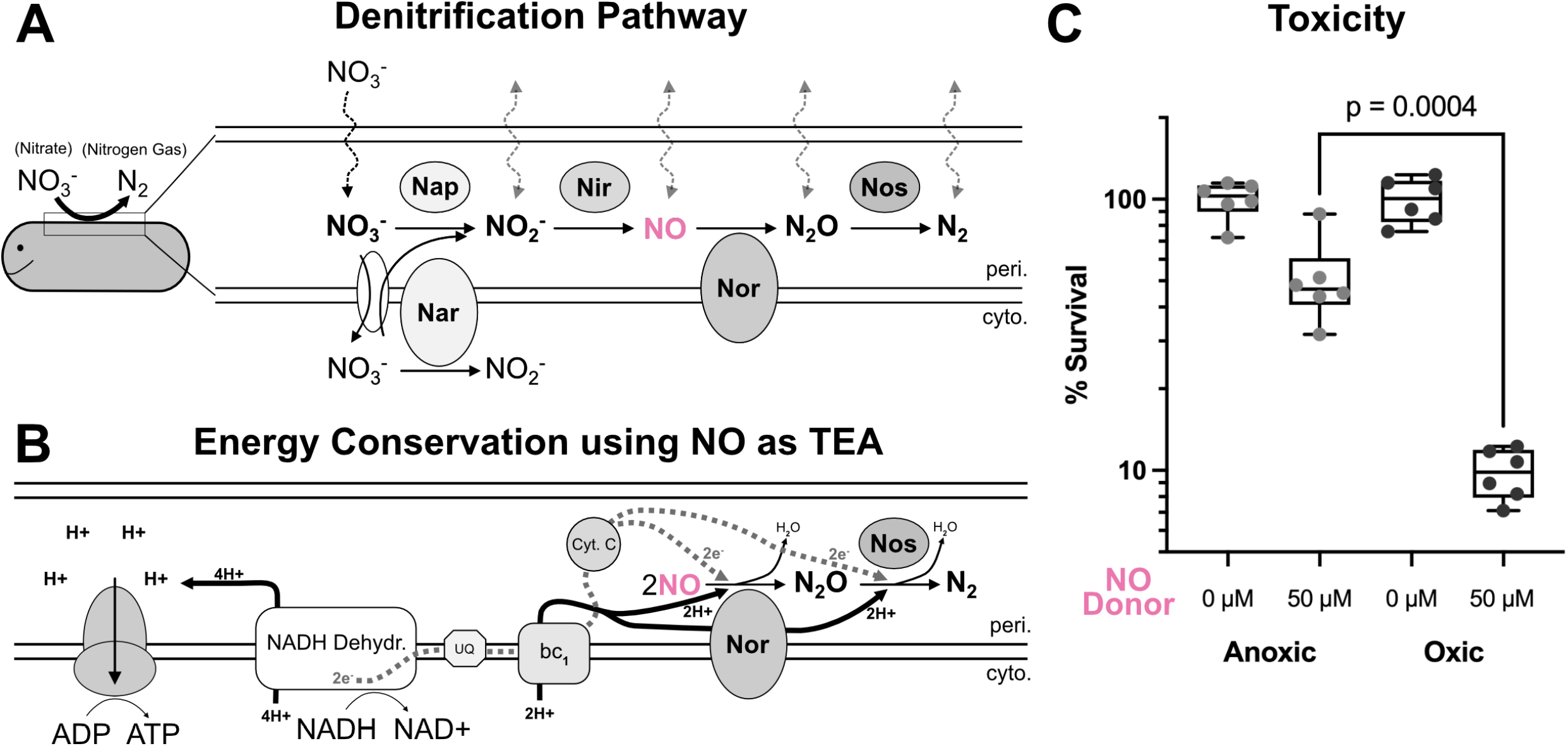
The agathokakological roles of nitric oxide in *Pseudomonas aeruginosa* growth. (**A**) WT PA14 sequentially reduces nitrate to nitrogen gas using five major enzymes. (**B**) Proposed mechanism of energy conservation using NO as the sole starting terminal electron acceptor (TEA). (**C**) WT PA14 survival after exposer to a short burst of an NO donor molecule under anoxic or oxic conditions. P value from unpaired t-test. Each dot represents one replicate, n=6 per condition.

To determine how the wild type (WT) PA14 strain responds to NO exposure, we incubated it with a common NO donor molecule called DEA NONOate under both anoxic and oxic conditions (Figure 1C). DEA NONOate releases 1.5 moles of NO per mole of parent molecule. WT cells were grown to mid-exponential phase, washed into anoxic buffer (pH 7.4) with and without a final concentration of 50μM NO donor (∼75 μM NO). This concentration was chosen to exceed that which has been shown to be tolerated by NO reductase (Nor) deficient mutants (∼15 μM) with the expectation that the WT strain could detoxify using its Nor ^16^. Cultures were incubated at room temperature in an anoxic glove box for one hour (donor half-life ∼16 minutes). After incubation, cultures were washed in fresh anoxic buffer and plated for colony forming units (CFUs) which revealed a 50% NO killing compared to an untreated control (Figure 1C). Next, we hypothesized that incubating cells with NO donor under oxic conditions would increase the toxic effects. After repeating this experiment under a normal atmosphere, NO exposure killing increased to 90% of the population (Figure 1C). These results suggest that NO as the sole TEA has the potential to contribute to energy conservation if certain conditions are met: that its concentration be maintained in the sub 75 μM range and that oxygen not be present in the microenvironment.

To generate NO at the microscale, we partitioned the PA14 denitrification pathway between two strains, a NO producer and a NO consumer, to make a synthetic community that could catalyze the complete denitrification reaction (Figure 2A). We reasoned that such a community would allow an NO producing strain to supply a low but steady concentration of NO to its partner consumer strain, minimizing toxicity issues and circumventing the need to work with NO donor molecules. Because of the sequential nature of the denitrification steps, we began by creating and characterizing several single, double, triple and quadruple clean deletion mutants to split the first and second half of the pathway at NO between two strains (Figure S1). After several iterations, we settled on a co-culture consisting of two strains that allowed us to observe NO cross-feeding via the presence or absence of growth under anoxic conditions (Figure 2A, co-culture 4 from Figure S1A). The NO producer strain, magenta outline, lacks both the NO reductase (Nor) as well as the N_2_O reductase (Nos). Compared to WT (black) under anoxic growth with nitrate (Figure 2B), the producer strain shows a dramatic defect in growth. This is consistent with previously characterized Nor mutants that can accumulate upwards of 15 μM NO before triggering a negative feedback loop that represses further nitrate and nitrite reduction, preventing further NO production ^16^. To ensure NO would be the sole TEA available for use, we designed the NO consumer strain to lack both the nitrate reductases (the proton pumping, membrane bound Nar and the redox balancing, periplasmic Nap), and nitrite reductase. Compared to WT, the consumer strain (cyan) also shows a dramatic growth defect, but unlike the producer strain, without NO inhibition, shows slight growth likely due to small amounts of carryover oxygen or built-up energy stores from the inoculum (Figure 2B). When these two strains are co-cultured to complete the pathway (dark blue), we see a large increase in total growth, as would be expected if the producer were relieved of deleterious NO build-up, and the consumer were provided with a low-level supply of an otherwise toxic TEA. (Figure 2B).

**Figure 2.**
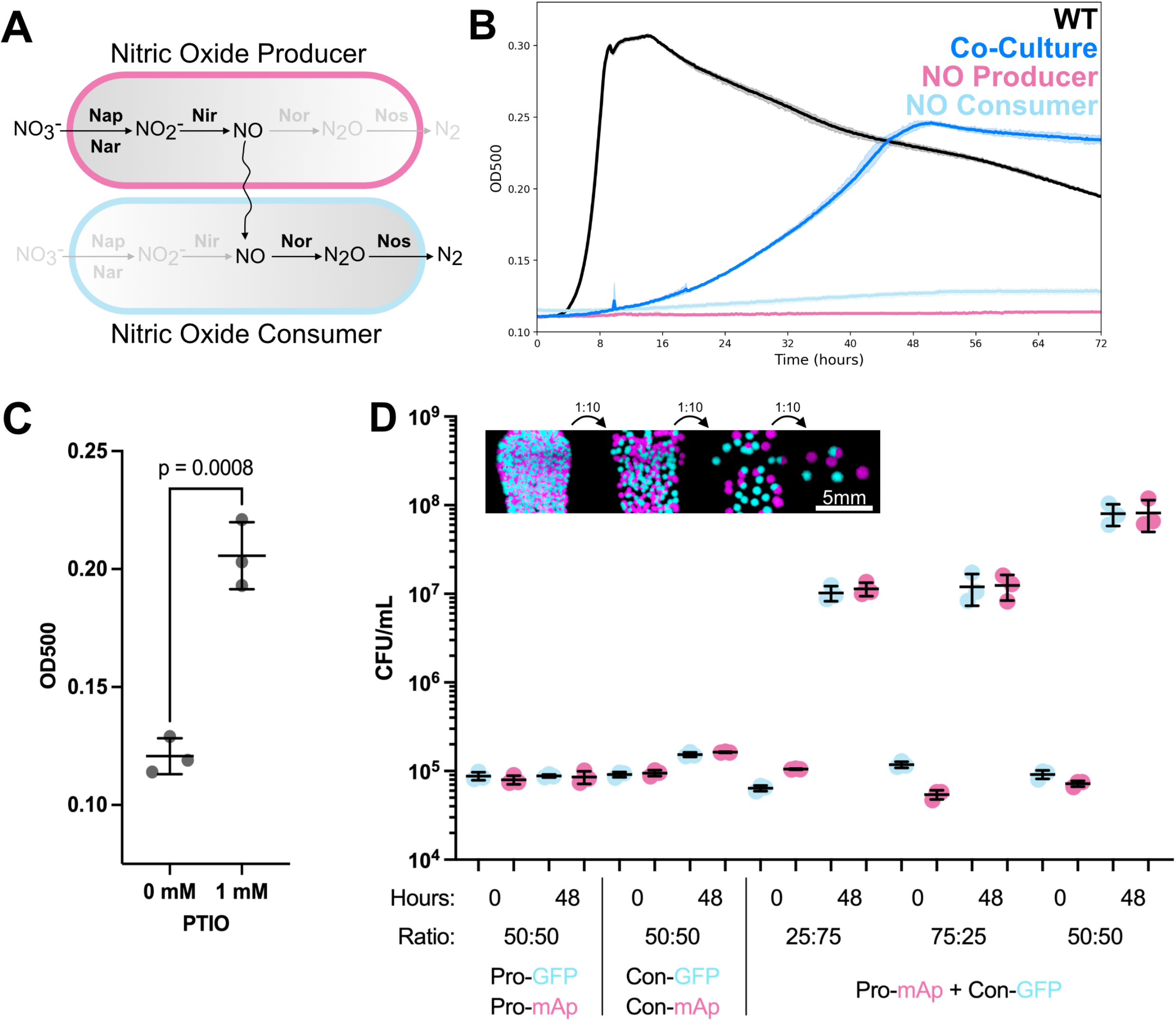
Design and characterization of synthetic nitric oxide cross-feeding community. (**A**) Schematic of NO producer (magenta) and consumer (cyan). Genes that have been cleanly deleted are grayed-out, and those remaining are shown in black. The pathway partitions where NO is produced and reduced. (**B**) Growth curve of strains in (A) grown individually and in co-culture (partitioned pathway), compared to WT (full pathway), over 72hrs under anoxic conditions with 5mM nitrate as sole provided terminal electron acceptor (TEA). Each growth curve represents n=4 replicates. (**C**) Growth rescue of producer strain using the chemical NO scavenger c-PTIO. Line and error bars represent means and standard deviations; P value from unpaired t-test. Each dot represents one replicate, n=3 per condition. (**D**) Different mixtures of the producer and consumer strains constitutively expressing either a GFP or mApple fluorescence reporter were grown as in (B) but at the beginning and end of a 48 hour incubation were plated for colony forming units. Consumer strain is Δ*narGHJI*Δ*nirS*Δ*napAB*Δ*ackApta*. Producer strain is Δ*norCB*Δ*nosZ*. See also Figure S1 and Table S1. Each dot represents one replicate, n=3 per condition

To verify that the growth of the strains in the anoxic co-culture was due to the removal of NO from the producer by the consumer, we functionally replaced the consumer with the chemical NO scavenger, c-PTIO, and grew the NO producer strain in its presence or absence. Compared to growth without the addition of c-PTIO restored NO producer growth, supporting the interpretation that NO reduction by the consumer drives growth in the co-culture (Figure 2C). Further, to address whether the consumer’s reduction of NO in the co-culture caused it to grow, we removed the NO reductase (Nor) from the consumer strain (Figure S1C). While this deletion did not alter the growth of this strain alone (Figure S1C, cyan), co-culturing this strain with the NO producer not only prevented, but also appeared to hinder co-culture growth completely (Figure S1C, dark blue). Finally, we asked if substrate level phosphorylation via pyruvate fermentation, a means of ATP generation for long term anaerobic survival in PA ^35^, might contribute to consumer growth by using NO in a redox balancing role as occurs for phenazines, another redox-active metabolite made by PA ^36^. As with Nor deletion, removal of genes responsible for substrate level phosphorylation (*ackA* and *pta*) did not have a measurable effect on maximum OD reached using a plate reader (Figure S1D, cyan). However, unlike Nor removal, the co-culture’s maximum OD was unaffected by the consumer’s ability to perform this process (Figure S1D, dark blue), suggesting consumer growth may be attributed to energy conservation from NO reduction as graphically depicted in Figure 1B.

These experiments suggest there is a net community benefit to strains in planktonic co-culture with a partitioned denitrification pathway under anoxic conditions compared to monoculture. To unpack this bulk phenotype and discern the effects of co-culture on the two members, we co-cultured the NO producer and consumer expressing constitutive fluorescence markers (GFP or mApple) for 48 hours. To track each strain’s growth in the mixed population, we performed serial dilutions of the anoxic culture and plated for CFUs under oxic conditions. Using a fluorescence microscope and image analysis, we calculated the change in starting cell numbers and ratio of the two strains at the final time point (Figure 2D). Alone, the NO producer, regardless of the fluorescence reporter used, showed no change in CFU after 48 hours (Figure 2D). This indicates NO production under these conditions reached inhibiting levels, but not necessarily enough to cause death. The NO consumer in the absence of the producer exhibited slight growth (Figure 2D), as seen in the growth curves (Figure 2B). Finally, we measured the growth dynamics of each strain in the co-culture. Regardless of the starting ratio (more producer, more consumer, or equal) both strains grew significantly, converging on an even ratio of producer to consumer by 48 hours (Figure 2D). These results demonstrate that NO, when supplied at a low but steady level, can serve as a TEA to support slow anoxic growth. This exchange promotes a mutualistic interaction where growth inhibition of the producer is alleviated by the consumer, which in turn gains a TEA; together, they flourish.

### NO cross-feeding under anoxic and sessile conditions is mutually beneficial and proximity dependent

Next, we wondered how spatial structure might influence the growth of the producer and consumer given that NO is a highly reactive and diffusible molecule. We hypothesized that, unlike under shaken planktonic growth conditions in which the whole population benefits, under spatially constrained anoxic conditions, proximity determines the success of community members. To test this prediction, we mixed fluorescently labeled cells of the same or differing genotypes in equal ratios, diluted the cultures, and sandwiched them between a coverslip and a nitrate-supplemented agar medium pad such that single cells were randomly distributed from one to dozens of cell lengths from each other. From these starting points, single cells could develop into aggregate biofilms over a 48-hour period. Figure 3A graphically represents the strain mixtures shown in Figure 3B as well as their predicted interactions based on the results of our planktonic growth experiments. As controls, we mixed WT with WT to explore how two cells with the full pathway and no growth defects would interact under these conditions, as well as the consumer and producer strains with themselves. Finally, the producer and consumer strains were mixed to visualize their interaction.

**Figure 3.**
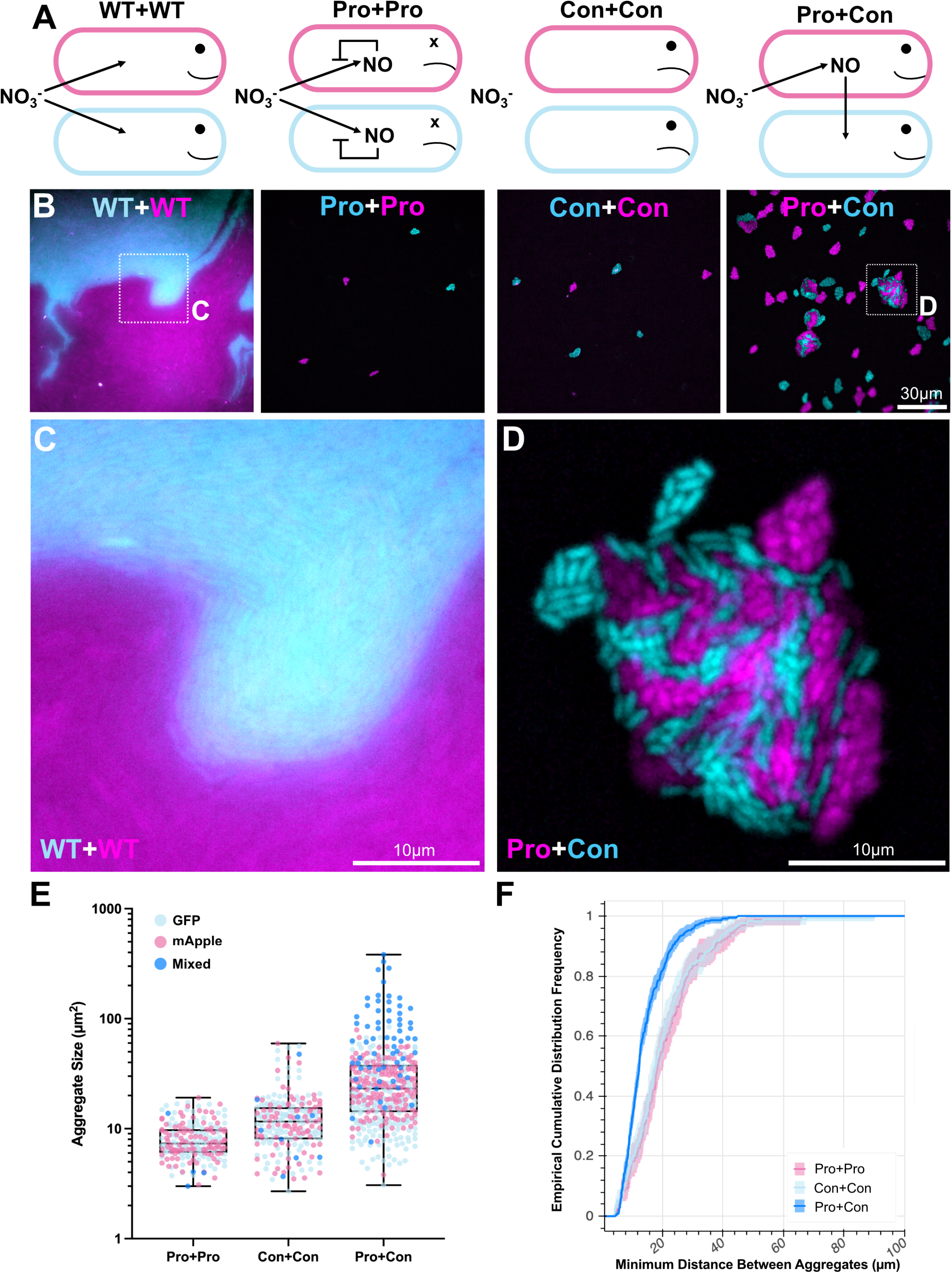
Patterning of producer and consumer strains under sessile anoxic conditions. (**A**) Phenotypic predictions. (**B**) Representative images showing growth patterns of indicated strain mixtures constitutively expressing either GFP or mApple after anoxic growth for 48hrs under an agar pad with nitrate provided as sole TEA. (**C**,**D**) Zoom-in of WT mixture showing strains growing as clonal patches (C) compared with zoom-in of the co-culture showing intermixing (D). (**E**) Quantification of each aggregates (dots) size and color-coded based on whether the aggregate contained just GFP, mApple or both (mixed) cells. (**F**) ECDF plot of the Euclidean distance between each aggregate and its closest neighboring aggregate. Consumer strain is Δ*narGHJI*Δ*nirS*Δ*napAB*Δ*ackApta*. Producer strain is Δ*norCB*Δ*nosZ*. E and F display values from at least 190 aggregates measured from at least 25 images per condition, pooled from two independent replicates. See also Figure S2.

Using widefield microscopy, we observed the two strains (GFP/mApple) in the WT mixture grew as large clonal expansions with little single cell intermixing, indicating neither was impacted by the other (Figure 3B-3C) ^37,38^. In contrast, both the Pro+Pro and Con+Con mixtures showed a dramatic reduction in aggregate size (proxy for total growth accumulation until observation, 48 hours) compared to WT (Figure 3B), in line with the inability of these strains to reduce nitrate, the supplied TEA. Interestingly, in order to observe this low baseline of growth for the consumer strain when grown with itself, it was necessary to use a consumer strain deficient in pyruvate fermentation. Whereas optical density was unable to resolve a difference between consumer strains +/- fermentation capability, this single cell imaging approach significantly increased our growth detection sensitivity, enabling measurement of subtle, but significant, differences (Figure S2A-2C, compare with Figure S1D, cyan). If we compare the WT (Figure 3C) and co-culture (Figure 3D) under these conditions, most strikingly we observe an increase in single cell intermixing between the two strains for the co-culture (Figure 3B vs 3D). The two WT strains (GFP/mApple) appear to grow as clonal expansions, pushing against one another rather than maximizing cell-cell contact (Figure 3C). Conversely, the co-culture shows slower growth compared to WT (same under planktonic growth), but appears to favor cell-cell contact between strains, forming mixed aggregates (Figure 3D), suggestive of a reliance of each partner on one another for continued growth ^39^. These mixed aggregates likely resulted from close starting positions of producer and consumer cells.

In addition to intermixed co-culture aggregates, we also observed a difference in aggregate size between single genotype mixtures and the co-culture (Figure 3E). If we compare producer aggregates (Pro+Pro, GFP or mApple dots) to the producer aggregates when grown with the consumer strain (Pro+Con, mApple dots) we see a significant increase in aggregate size (Figure S2D); thus, even when the producer is not in direct contact with the consumer, it appears to benefit from NO removal at distance. The consumer strain (Con+Con, GFP or mApple dots) also showed a small but statistically significant increase in size even when not in direct contact with the producer (Pro+Con, Cyan dots) (Figure S2D). However, the largest aggregates were those that contained a mixed population of producer and consumer strains (blue dots) (Figure 3E and Figure S2D).

Importantly, the increase in aggregate size even when the co-culture strains where not in direct contact indicates that while single cell intermixing can occur when the two strains are seeded at close initial starting positions (Figure 3D), without a proximal sink, NO can diffuse outward from producer cells to reach consumer cells thereby increasing the interaction range. NO’s ability to rapidly traverse the distances separating source from sink cells in our mixed cultures is not surprising given prior results showing that NO can diffuse over 100 μm within 1 second ^40^. To determine whether diffusion alone could explain our results, we measured the Euclidean distance between the centers of each aggregate at 48 hours in producer-alone, consumer-alone and co-cultures. When we compare the minimum distance between each aggregate and its closest neighbor, we find that all are within 100 μm of each other, with the majority between 20-60 μm (Figure 3F). Notably, there is a slight decrease in distances between co-cultured aggregates, likely due to better growth given the exchange of NO. Together, these results extend our planktonic growth observations into a spatial context, revealing how community members with a partitioned denitrification pathway can interact under anoxic conditions where NO can freely diffuse.

### Oxygen availability enhances NO toxicity, thereby changing the nature of the co-culture interaction

Recognizing that oxygen presence increases NO toxicity (Figure 1B) and that many environments where denitrification occurs experience dynamic changes in oxygen and nitrate consumption coupled to increased production of nitric and nitrous oxide ^5,41^, we wondered what would happen to the NO producer strain and co-culture when grown in a heterogeneously oxygenated environment. Strains such as PA14 are facultative anaerobes, meaning they preferentially respire oxygen when it is available (oxic conditions), but when oxygen is limited, they induce pathways (such as denitrification) that permit energy conservation using alternative TEAs. A classic example of this phenomenon is when cells grow planktonically but are not shaken during growth. Cells at the surface of the medium reduce oxygen faster than it can diffuse into the culture, resulting in an oxygen gradient (Figure 4A, left) ^42^. To expand our understanding of when, where and how NO-cross feeding operates, we exploited this phenomenon and measured growth in shaking or standing “oxic” cultures. We hypothesized that strains harboring a partitioned denitrification pathway would exhibit growth that depends on the amount of oxygen and nitrate present, in a manner reflecting the agathokakological effects of oxygen and NO on cells.

**Figure 4.**
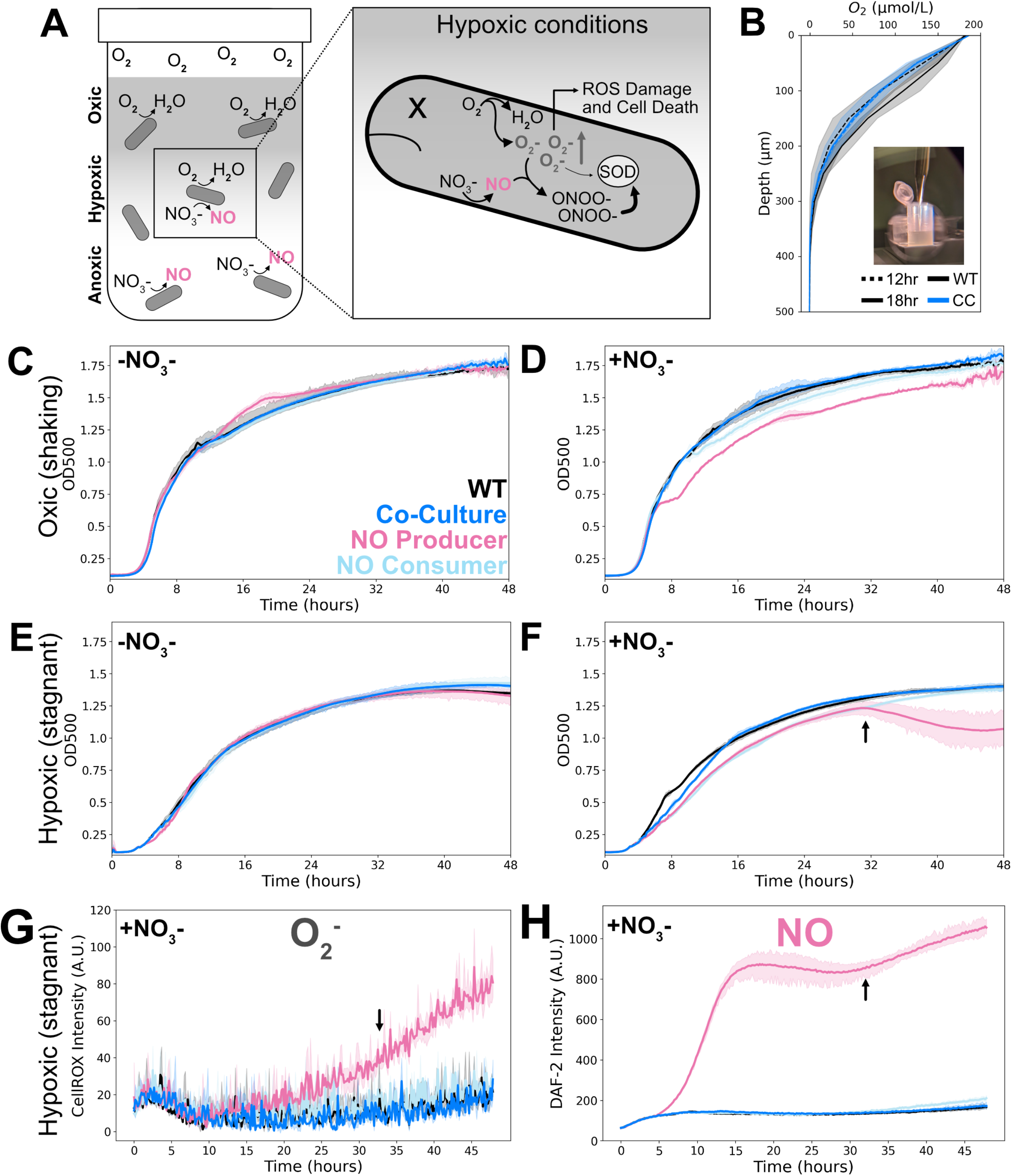
Oxic vs hypoxic growth of planktonic cultures supplemented with and without nitrate. (**A**) Graphical prediction of the heterogeneously oxygenated environment created by stagnate liquid culture incubation (left). Expected consequences of the NO producer growing under this environment (right). Superoxide (O_2_^-^) produced during aerobic respiration is kept below toxic levels by superoxide dismutase (SOD). The byproduct of NO and O_2_^-^, ONOO^-^, can out-compete O_2_^-^ for SOD causing an accumulation of O_2_^-.^ (**B**) Oxygen profiles from non-shaking cultures of the WT compared to profiles from the co-culture at two time points to confirm heterogeneity. Each trace represents n=3 replicates. (**C-F**) 48hr growth curves cultured under the indicated conditions. Evenly oxygenated environments (“oxic-shaking”) were achieved by shaking cultures during incubation (C,D), while unevenly oxygenated environments (“hypoxic-stagnant”) were achieved by culturing without shaking (E,F). (**G-H**) Fluorescent dye readings of superoxide (G) and nitric oxide (H) accumulation over a hypoxic growth curve with nitrate (as in F). Consumer strain is Δ*narGHJI*Δ*nirS*Δ*napAB*. Producer strain is Δ*norCB*Δ*nosZ*. Each trace in C-H represents n=3 replicates. See also Figure S3.

To test this hypothesis, we first observed how stagnantly grown cultures created an oxygen gradient by measuring the oxycline of both the WT and co-culture at multiple time points using an oxygen microelectrode (Figure 4B). The oxycline was present and consistent between strains and time points. As a baseline, we tracked the growth of the producer and consumer strains, alone and in co-culture, over 48-hours in a continuously shaking plate reader to prevent an oxygen gradient from forming. In the absence of nitrate all strains grew similarly, as expected, indicating the genetic modifications had no off-target effects (Figure 4C). Conversely, with 5 mM nitrate added to the medium, the consumer showed a slight growth defect compared to the WT and the producer displayed a brief growth arrest midway through exponential phase, with growth then paralleling that seen for the WT or mixed producer + consumer cultures (Figure 4D). Despite continuous shaking, at these cell densities, we surmise from prior work that these strains began to experience oxygen-limitation during mid exponential phase ^42^.

To further induce hypoxia, we performed the same planktonic growth experiment without the plate reader shaking, mimicking the conditions leading to the oxycline shown in Figure 4B. Again, in the absence of nitrate, we did not see a difference between the strains (Figure 4E). However, in the presence of nitrate, all but WT showed a slight lag in growth (Figure 4F). Importantly, even without the ability to reduce nitrate, the consumer was able to eventually reach the same final OD as WT. Conversely, the producer strain showed a dramatic decrease in OD later in the growth curve, suggestive of cell lysis. To check whether this reduction in OD was due to a reduction in viable cells, we counted CFUs. Compared to WT viability at the 48hr time point, there was a significant reduction in producer CFU (Figure S3A). To confirm whether growth and prevention of cell death in the co-culture was caused by NO consumption, we repeated the growth curve but this time with a consumer strain lacking the NO reductase (Nor) (Figure S3B). When the Nor deficient^-^consumer strain was co-cultured with the producer under hypoxic conditions, we observed a reduction in OD similar in timing and amount to the producer growth alone. Because the consumer alone reaches the same final OD as WT, yet would otherwise die if it lacked NO reduction ability in the co-culture, we propose that under hyp(oxic) conditions the consumer plays a beneficial role in the mixed community

Finally, to assess whether concomitant aerobic respiration and denitrification were occurring under these conditions, we tracked the accumulation of superoxide (O_2_^-^) and NO. O_2_^-^ is a by-product and intermediate of oxygen respiration that is typically removed by superoxide dismutase (SOD) (Figure 4A, right). However, NO readily reacts with superoxide (O_2_^-^) to form the reactive nitrogen species, peroxynitrite (ONOO^-^), which can out-compete superoxide for SOD leading to an accumulation of reactive oxygen species and lead to oxidative stress (Figure 4A, right) ^17^. Thus, we reasoned that the increase in cell death observed under hypoxic conditions for the NO producer might correlate with an inability to scavenge superoxide. To test this idea, we used a O_2_^-^ sensitive dye (CellROX, Figure 4G) and an intracellular NO sensitive dye (DAF-2DA, Figure 4H) to detect their production. Both O_2_^-^ and NO increased over the growth curve in the producer strain when grown alone (Figure 4G-4H), with a rise in signal around 30 hours, correlating with the drop in OD (Figure 4F, arrows). This decrease in OD and increase in O_2_^-^ signal likely occurs due to cell death and thus a collapse of the oxygen gradient leading to a cascade of further ROS production.

### NO cross-feeding under (hyp)oxic and sessile conditions leads to spatial behavioral patterns

Unlike under anoxic conditions where the consumer strain gains a TEA that allows it to grow, we wondered what role its reduction of NO would play under heterogeneously oxygenated, sessile growth conditions. To achieve this, we grew diluted cells on agar pads under oxic conditions into aggregate biofilms (Figure 5A). Unlike the anoxic experiments, here cells initially had full access to oxygen and thus we expected them to behave as in planktonic oxic growth conditions. Over time, we expected cell density to rise and oxygen to diminish, prompting greater nitrate reduction to NO by the producer strain, as seen under planktonic hypoxic growth conditions. The question became: how would the fixed organization of NO producers and consumers in space influence their behavior over time?

**Figure 5.**
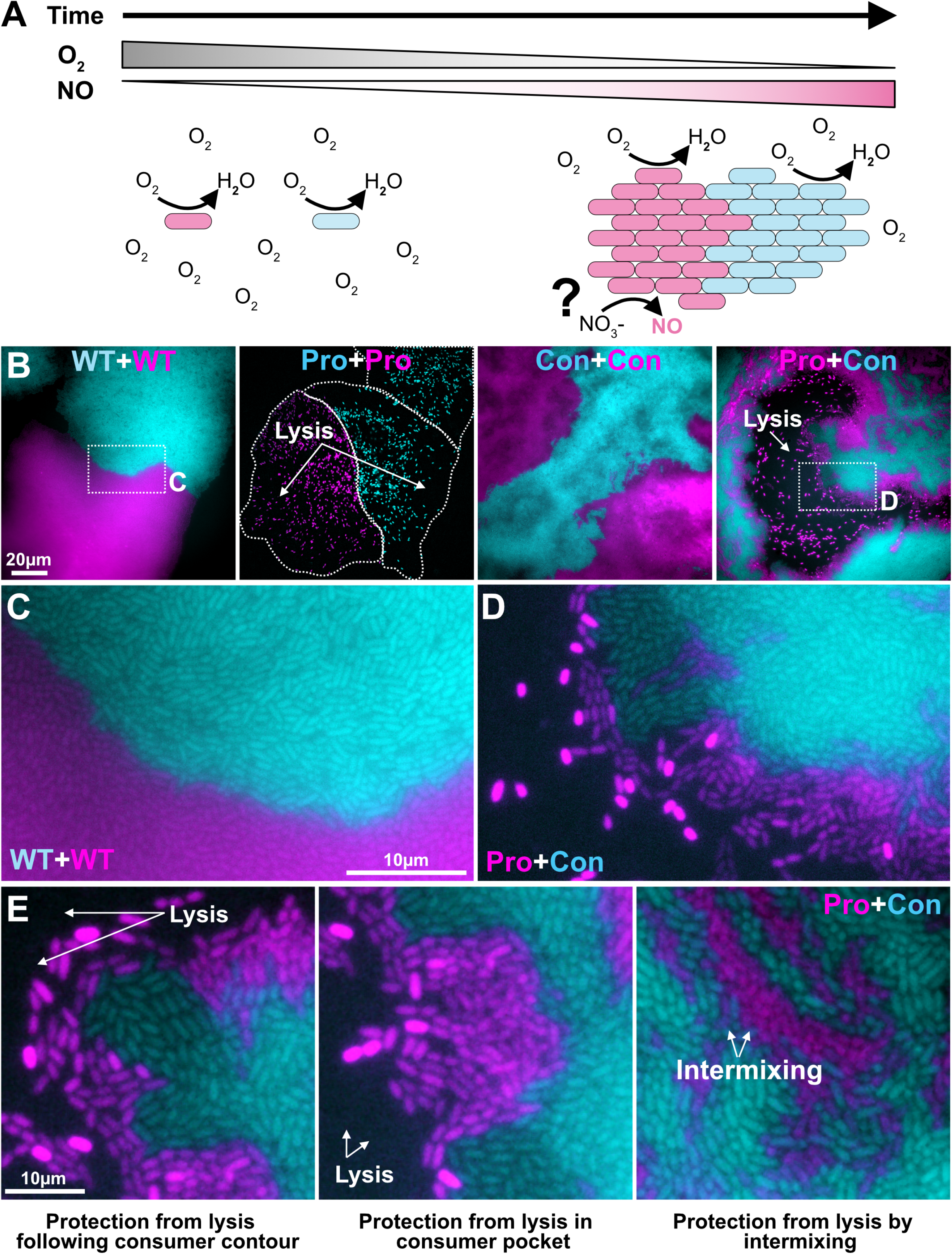
Patterning of producer and consumer strains under sessile oxic conditions. (**A**) Graphical prediction for NO producer and consumer strains growing initially in the presence of replete oxygen. Overtime oxygen consumption will lead to hypoxia and presumably a higher and higher production of NO. (**B**) Representative images of indicated strains grown as biofilms for 24hrs under oxic conditions in the presence of nitrate. (**C**) Zoom-in of WT clonal expansion showing no intermixing. (**D**) Zoom-in showing producer survival adjacent to the consumer expansion interface; note that cells appear brightest prior to lysis. (**E**) Three additional high magnification callouts to spotlight specific structural features. Consumer strain is Δ*narGHJI*Δ*nirS*Δ*napAB*. Producer strain is Δ*norCB*Δ*nosZ*. See also Figures S3 and S4.

To visualize interactions between NO producers and consumers, we mixed the strains with themselves or each other and incubated them under oxic conditions on nutrient agar pads containing nitrate. After 24 hours of growth, we observed with widefield fluorescence microscopy that WT, like under anoxic conditions, grew into aggregate biofilms; however, rather than monolayers, these biofilms appeared thicker perhaps due to their access to oxygen and ability to support both oxic and anoxic populations (Figure 5B). Strikingly, the producer only mixture (Pro+Pro) also showed clonal patches (outlined with a dotted white line, Figure 5B) but with very few cells remaining fluorescent. To investigate further, we examined fluorescence and corresponding phase channels more closely (Figure S3C-3E). Cells that maintained their fluorescence after this incubation period also maintained their opacity in the phase channel. Cells no-longer showing fluorescence appeared transparent, leading us to conclude that they have undergone cell death and lysis as seen under hypoxic planktonic growth (Figure 4F). Conversely, the consumer only mixture (Con+Con) grew into clonal patches, (note much larger than under anoxic conditions due access to oxygen as a TEA) though they did not appear as dense as WT owing to their inability to support anoxic growth with nitrate.

The co-culture, however, provided several new insights. First, there was an increase in single cell intermixing (compare Figure 5C (no mixing) and 5D (mixing)). Second, unlike under planktonic growth conditions where the whole co-culture population benefits from interacting, here there appeared to still be lysis of producer cells when not in proximity to consumer cells (Figure 5D, Figure S4A shows lysis overtime). This indicates that spatiotemporal, microenvironmental changes induce metabolic dynamics that unlock a spatial niche where cross-feeding provides a selective advantage as protection from lysis for the producer strain. Such protection spatially patterns in several ways (Figure 5E): producer cells follow the consumer contour; others appear to be protected by a pocket formed by consumer cells; and still others are protected in regions of high intermixing. Knowing that the producer cells would lyse without local NO depletion under hypoxic conditions (Figure 4F) permits us to link such cellular spatial arrangement to physiological activity.

### NO-dependent, co-culture interaction patterns dynamically reflect changes in the microenvironment

To test our predictions about how these spatial patterns arise, we repeated agar pad imaging as in Figure 5, but varied whether or not the consumer strain encoded the nitric oxide reductase (Nor). First, we asked whether the presence of Nor decrease ROS stress by preventing the downstream generation of superoxide. To address this, we included CellROX in the agar from the beginning of the experiment to visualize super oxide accumulation using confocal microscopy (Figure 6A-6D). The consumer channel was used to generate a mask and measure CellROX fluorescence intensity of the consumer only space (illustrated in Figure S4B). Comparing the results of consumer strains with (Figure 6A) and without (Figure 6B) Nor, we found a dramatic increase in CellROX staining at the border between the strains, and a measurable increase in the consumer cells surrounding the producer cells. This suggests NO diffusion leads to ROS generation in the absence of NO reduction by the consumer. Next, we asked how the consumer affects the producer cells in these biofilms by measuring the remaining producer cell area as a function of available, non-consumer space (inverse of consumer mask, illustrated in Figure S4B). The results show a dramatic reduction in producer area when the consumer lacks the ability to reduce NO (Figure 6D).

**Figure 6.**
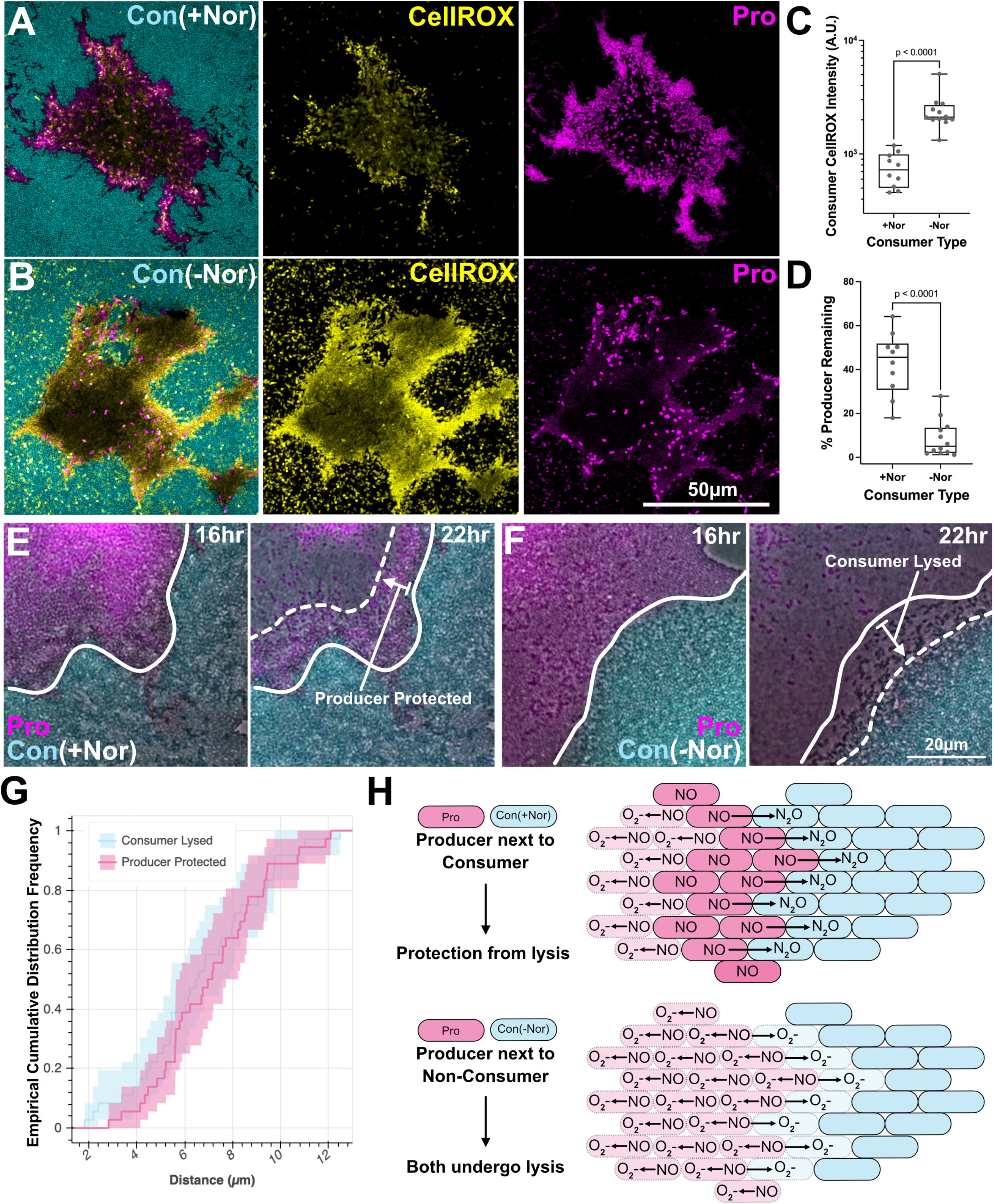
Dynamic spatial patterning of NO partitioned community. **A-D**) Visualization and quantification of spatial superoxide production and lysis as a function of NO consumers presence (+Nor) or absence (-Nor) of the NO reductase. Specified strains were grown as in Figure 5 but including the superoxide sensitive dye CellROX (Yellow). (**C**) Mean CellROX intensity of consumer cells was measured whether it contained the NO reductase (A) or not (B). (**D**) Using the inverse of the consumer region as a pre-lysis producer area mask compared to the remaining producer area (unlysed), a % remaining metric was obtained. P values from unpaired t-tests in C and D. Each dot represents measurements from single images (n ≥ 10). (**E-F**) To measure lysis range in the presence or absence of NO reductase, time lapse imaging was performed from pre- and post-lysis. (**G**) ECDF of lysis/protection from lysis range was measured from 12 regions each from three aggregate co-culture interfaces (between solid and dashed lines as drawn in E and F, n=36). (**H**) Proposed model in which the NO producer and consumer strains both grow clonally until the microenvironment changes such that the producer begins to produce NO and lyse. The consumer acts as a sink for NO in close proximity to the producer, relieving NO toxicity and preventing the producers and own lysis. Consumer strain is Δ*narGHJI*Δ*nirS*Δ*napAB*. Producer strain is Δ*norCB*Δ*nosZ*. See also Figure S4.

Finally, these end time point measurements assumed where the border between consumer and producer started. The increase of CellROX staining at the border when the consumer lacked NO reductase suggests it could be undergoing lysis itself. This led us to ask if the consumer lysis when NO cannot be reduced matches the protection afforded to the producer when it can be reduced. We performed time lapse microscopy of the two co-cultures (consumer +/- Nor) both pre- and post-lysis. This allowed us to mark the initial border (Figure 6E, solid line) between the strains and draw a line indicating the extent of protection (Con + Nor) or induction (Con - Nor) of lysis (Figure 6E, dashed line). Indeed, the consumer strains without Nor undergo lysis at the border with the producer (Figure 6F), suggesting the major role for its NO reduction under these conditions is first to protect itself from toxicity, with a secondary benefit for the producer. Quantifying this phenomenon, we measured the range of protection (distance) conferred to the producer by the consumer (arrows, Figure 6E; pink line-Figure 6G). When the consumer lacks NO reduction capacity, all of the producer is vulnerable to lysis and the equivalent protection range manifests as a lysis range in the consumer population due to NO diffusion (Figure 6F, cyan line-Figure 6G). Overall, these ranges (2-12 μm) are much shorter than the interaction distances observed under anoxic growth conditions (20-60 μm, Figure 3F). We interpret this to be a consequence of NO’s high reactivity with oxygen effectively shortening its diffusive range. Figure 6H graphically depicts our working model, showing how initially non-interacting cells can develop a high stakes interaction with respect to fitness as the microenvironment changes.

## Discussion

Awareness of the diversity of microbial metabolic interactions is expanding with improved techniques for identifying new pathways and observing interactions ^1,43–47^. In the face of this progress, gaining a better understanding of how these interactions may shift as a function of dynamic oxygen gradients is an important goal. We developed a synthetic microbial community linked by the exchange of NO, a redox active molecule whose biological effects are tuned by the presence of oxygen, as a step towards this end. Our work shows that the interactions that drive NO intercellular exchange and reduction are induced and sensitive to the genetic content of community members and microenvironmental gradients, knowledge of which leads to predictable organizational outcomes.

Given denitrification may be a community effort ^23^, and that each community member may contain a different suite of relevant enzymes, our studies with a synthetic community suggest that certain agathokakological intermediates, such as NO, may play an outsized role in structuring natural microbial communities. Linking environmentally-dependent metabolic interactions to microscale microbial organization is an important goal ^48–53^. Our findings spotlight dynamic changes in local oxygen for its ability to tune both the nature and need for a cross-feeding relationship. In our example case, NO can generate toxic byproducts under oxic conditions, leading to cellular growth arrest. Yet under anoxic conditions, NO can be a lifeline, serving as a TEA and promoting energy conservation and growth. These contrasting effects are important because oxygen exhibits steep environmental gradients over small spatial scales ^5,54^. Our work draws attention to the fact that oxygen gradients can shape microbial interactions both in planktonic and sessile communities, with the scales over which these interactions occur being set by the proximity of interacting cells and their local microenvironment, which they in turn alter in space and time.

An important next step will be to determine whether mixtures of natural isolates with partitioned metabolic pathways play by the same rules. Our work gives rise to testable predictions about when and where NO cross-feeding may occur. For example, it is well appreciated that NO is an important intermediate species in the nitrogen cycle ^22^, whose turnover rates may decide the fate of soil nitrogen stores. Interestingly, recent metagenomic studies indicate that within certain habitats, there an enrichment for organisms that can only reduce NO to N_2_O, an important greenhouse gas ^45^. Thus, an understanding of what constrains NO reduction may provide insight into a metabolic process with important environmental consequences. Recent studies have drawn attention to the potential for partitioned pathways to influence the rate of nitrogen flux in soils by resident microbes ^23,45^. Our work attempts to directly link structure and function of such partitions as is suggested by the genomic content of environmental isolates ^45^. In an entirely different context, our finding that *P. aeruginosa* can thrive by reducing NO provided by another cell lends credence to the hypothesis that PA may survive the host immune NO attack by its reduction to N_2_O, and may correlate with the appearance of N_2_O in the breath gas of cystic fibrosis patients ^54^. Importantly, *P. aeruginosa* can be found proximal to host immune cells in these infected environments ^44^.

Finally, we note that NO is only one of many redox active molecules (RAMs), both organic and inorganic, that may help structure microbial communities over small spatial scales as a function of oxygen availability. For example, other RAMs such as phenazines can function as natural toxins or beneficial agents of nutrient acquisition and/or redox homeostasis, depending on the context ^9^. Similarly, sulfide has long been known to play nuanced roles in nature, with its concentration-dependent toxicity influencing which phototrophic species are able to thrive ^55^ and where different energy-conserving processes can occur within microbial mats ^56,57^. Many other parameters may be subject to similar dynamic spatiotemporal gradients, such as carbon, pH and light, and are likely to also play important roles in determining the niches where cross-feeding or other types of metabolic interactions can occur (^29,30,58^). Towards the goal of gaining a predictive understanding of why diverse microbial communities are structured the way they are, a constructive next step will be to model these interactions in light of fluctuating environmental parameters, iterating with experiments until cross-cutting principles can be identified with predictive power.

## Acknowledgments

The work was supported by the National Institutes of Health (R01HL152190 to D.K.N.) and the National Science Foundation Graduate Student Fellowships Program (to S.A.W.). We would like to thank the current and past Newman Lab members for feedback and discussions. Specifically, we thank Drs. Darcy McRose and Avi Flamholz for manuscript review, Dr. Melanie Spero for molecular technique instruction, Drs. Reinaldo Alcade, Georgia Squyres, and Zachary Lonergan for fruitful discussion and assay troubleshooting, and Dr. Michael Piacentino for help with programming and encouragement. Some of the imaging was performed in the Biological Imaging Facility of the California Institute of Technology, with the support of the Caltech Beckman Institute and the Arnold and Mabel Beckman Foundation.

## Author contributions

SAW and DKN contributed to study conceptualization, data interpretation and writing – review and editing. SAW developed methodology, performed experiments, data curation, and original manuscript draft.

## Inclusion and diversity

One or more of the authors of this paper self-identifies as a member of the LGBTQIA+ community. We support inclusive, diverse, and equitable conduct of research.

## STAR Methods

### KEY RESOURCES TABLE

See attached table

### RESOURCE AVAILABILITY

#### Lead Contact

The lead contact of this study is Dianne Newman who will fulfill all requests for additional information, resources, and reagents.

#### Materials availability

Bacterial strains and constructs that were generated in this study are available upon request to the lead contact.

#### Data and code availability

This paper does not report original code. All source images, raw data, and any other information necessary to replicate these analyses will be shared by the lead contact upon request.

### EXPERIMENTAL MODEL AND SUBJECT DETAILS

#### Bacterial strains

All strains used are clean deletion mutants originating from the wild-type (WT) *Pseudomonas aeruginosa* UCBPP-PA14 strain as described below in the Method Details. Strains were stored as 50% glycerol stocks at -80°C.

#### Bacterial growth conditions

Before use, strains were streaked from frozen stock onto Bacto agar (BD, Sigma) plates containing LB (Miller, BD, Sigma) or LB supplemented with 50 μg/mL gentamicin (GoldBio) to select for retention of expression cassettes within fluorescent strains, and grown overnight at 37°C. From streak plates, a dab of culture was used to inoculate 5 mL liquid cultures in LB or LB plus 50 μg/mL gentamicin liquid media. Cultures were shaken at 250 RPM at 37°C overnight until fully saturated at stationary phase. To standardize inoculum sizes between strains, each strain’s optical density (OD_500nm_) was measured by spectrophotometry (Beckman Coulter). Strains were then each washed and diluted to an OD of 1 via centrifugation and resuspension in phosphate buffered saline (PBS). Strains were then either further diluted directly, or first mixed at equal ratios for co-culture experiments and then diluted to final experimental concentrations. For all experiments, a Low Salt LB (LSLB) was used instead of standard LB powder due to microelectrode reactivity to the high NaCl concentration. The composition of LSLB was 141 mM Tryptone (BD, Sigma), 16mM Yeast Extract (BD, Sigma), 45mM NaCl (Fisher Chemical), which were dissolved in Mili-Q water and autoclaved. Nitrate was supplemented into media where indicated by diluting from a 1M stock of KNO_3_ (Fisher Chemical).

### METHOD DETAILS

#### Construction of mutant bacterial strains

Table S1 contains a full list of primers used in the creation of deletion constructs and mutant strains used in this study. Briefly, a 1kb fragment up- and downstream of the target gene was amplified via PCR and cloned into the pMQ30^59^ plasmid using Gibson cloning (NEB) and transformed into *E. coli* cells as previously described ^60^. Deletion constructs were then introduced to parent strains (WT PA14 and or deletion strains of PA14, see Table S1) via triparental conjugation. *E. coli* plasmid and helper strains were selected against by bead plating on VBMM plates containing 50 μg/mL gentamicin ^61^. Roughly 50% WT and 50% clean deletion colonies were then obtained by bead plated on a 10% sucrose plate to induce homologous recombination with the construct containing the regions up- and downstream, but lacking the targeted gene locus ^62^. Clean deletion strains were confirmed via PCR genotyping using primers that span the deletion region and sequencing of the amplified product. Finally, anaerobic growth challenges revealed expected physiological effects (e.g. ΔnirS was unable to grow on nitrite, etc). Fluorescence strains were created as previously described using plasmids driving GFP/mApple expression under control of the constitutive ribosomal promoter rpsG introduced into the Tn7 site of specified strains using tetra-parental conjugation^62,63^. *E. coli* helper strains were again selected against using VBMM containing 50 μg/mL gentamicin. Confirmation of insertion was performed via microscopy, visualizing colonies in both green and red fluorescence channels, and selecting those that were bright in their respective channel above background autofluorescence.

#### Serial dilution and colony forming unit (CFU) counting

Liquid cultures were plated for colony forming units and back calculated to assess number of cells. Typically, 100 μL of culture was placed into the well of a 96 well plate (at least three technical replicates were assessed). 20 μL of culture was then diluted seven times in serial 1:10 dilutions in PBS. 10 μL of the dilutions were then plated on a LB agar plate using a multichannel pipette. The plates were then slanted vertically and tapped to allow the 10 μL to spread in a line. All dilution plates were incubated under normal atmospheric conditions, in an incubator at set to 37°C for roughly 24 hours. After incubation, colony counting was performed either manually or with an imaging and image analysis pipeline (when fluorescence imaging was used, see below). Dilutions with 100-200 colonies were used to calculate colony form units (CFU/mL) by multiplying this number by the dilution factor.

#### Growth curves

Anoxic growth curve media was placed in anaerobic chamber 72 hours prior to experiment to allow time for oxygen to degas from media. 1 OD_500nm_ cultures were brought into chamber and further diluted to 0.005 OD_500nm_ in anoxic media. Growth curves were performed using a BioTek Synergy 4 plate reader stored in an anoxic chamber and set to 37°C with shaking and measured OD_500nm_ every 10 minutes. Oxic (shaking) and Hypoxic (stationary) growth curves were set up as above but under normal atmosphere. Oxic growth curves were performed using a Tecan Spark 10M set to orbital shaking. Hypoxic growth curves were performed using a Spectramax M3 (Molecular Devices) plate reader set to not shake. Each condition was performed in technical triplicate or quadruplicate. Each well contained 200 μL of culture with 40 μL of autoclaved mineral oil added to the top of each well to prevent evaporation of 48-72-hour growth curves. Traces from growth curves were analyzed and presented by plotting mean OD_500nm_ over time (dark center line) and 95% confidence intervals from technical replicates (shaded area) using Seaborn^64^ plotting packages in Python ^65^.

#### Nitric oxide and superoxide quantification

Levels of nitric oxide and superoxide were quantitated using fluorescent dyes during hypoxic growth curves. NO was quantitated using 50 μM DAF-2DA (Calbiochem, Sigma) which is internalized by cells where esterases free DAF-2DA to interact with NO producing a green fluorescence signal ^66^. Superoxide was quantitated using 5 μM CellROX Deep Red (Thermo), which is non-fluorescent in a reduced state but exhibits a strong induction of fluorescence upon oxidation with reactive oxygen species ^67^. The Spectramax plate reader was configured to measure green fluorescence (Ex. 488 nm, Em. 520 nm, DAF-2) and far-red fluorescence (Ex. 640 nm, Em. 665 nm, CellROX). For pad imaging assays, CellROX was supplemented to the agar pads directly at a final concentration of 10 μM and imaged after significant biofilm growth had occurred to reveal superoxide localization. DAF-2DA was also used to visualize NO; however, due to weak single-cell signal and strong background autofluorescence in the green channel, we were unable to visualize the signal convincingly.

#### Oxygen profiling

To confirm the presence of oxygen gradients in standing cultures, we employed a 25 μm diameter oxygen microelectrode (Unisense), which was calibrated as previously described ^60^. Briefly, LSLB media incubated at 37°C was used to calculate the high point O_2_ concentration. A zero-point calibration was obtained by dissolving 2 g Sodium Ascorbate in 100 mL 0.1 NaOH, which scavenges O_2_ from the medium. Using the Unisense Sensor Suite Software to save calibration values and control a motorized micromanipulator for profiling, O_2_ was measured from the surface of hypoxically incubated planktonic cultures in 1 mL Eppendorf tubes. The surface was set by eye and confirmed by a reduction in mV signal. O_2_ was then measured over a 1 mm distance below the surface at 50 μm intervals. Two measurements were taken at each position, and three technical replicates were averaged and displayed as the mean value with 95% confidence intervals (Figure 4B). Cultures were prepared as in growth curves but maintained as 1 mL volumes in tubes rather than aliquoting into 96-well plates. Mineral oil was not added due to potential incompatibility with electrodes.

#### Agar pad imaging assay

We determined that 1 OD_500nm_ cultures diluted to a starting concentration of 0.001 OD_500nm_ was sufficient to obtain single cells randomly distributed under agar pads. Agar pads were made by combining LSLB media with 2% w/v Noble Agar (BD, Sigma). Prior to use, the agar was melted by microwaving. To make equally sized agar pads, 100 μL was pipetted into each of several square molds (6-7 mm X 7 mm X 1.6 mm Depth ID, 25 mm X 75 mm, Grace Bio Labs), placed between two glass microscope slides, and allowed to solidify at 4°C until use. Anoxic pads were made 72 hours in advance and placed in an anaerobic chamber to degas. 5 μL of 0.001 OD_500nm_ dilution culture was added to the well of an 8-well LabTek dish (Thermo). Agar pads were then placed on overtop. Dishes were incubated at 37°C for 24 hours (oxic) or 48 hours (anoxic) in a sealed container (6.7 oz, Systema). When additional humidity was added, instead of distinct aggregate biofilms, cells appeared as under intermixed planktonic conditions; thus, additional humidity was not added. Time lapse imaging was performed incubating pads as described above, but just before lysis would occur (∼18 hours), dishes were sealed with parafilm and placed in the heated microscope incubation chamber for imaging. Anoxically-incubated cells were not initially fluorescent due to the inability for fluorophores to fold in the absence of oxygen. To overcome this, pads were placed at 4ºC without lids for one to two hours prior to imaging resulting in sufficient signal recovery to differentiate GFP and mApple positive cells.

#### Microscopy

All imaging was performed either using a Nikon Ti2 widefield fluorescence microscope or a Zeiss LSM800 microscope. End time point agar pad images were collected using the Nikon with a Plan Fluor 100x/1.30 oil phase objective and time lapse images were collected using a Plan Apo 40x/0.90 dry phase objective. GFP channels were collected using a 488 nm LED excitation light source and emission was collected with a FITC filter. The mApple channel used 561 nm LED light source and emission was collected with a TRITC filter. For time lapse imaging, an OKO Labs cage incubation unit was set to 37°C several hours before imaging to equilibrate the microscope stand and used for the duration. Images were collected every 20 minutes from multiple stage positions. Time indicated in figures represents time since the pads were inoculated. Both fluorescence CFU counting images from Figure 2D and the CellROX/Producer mass quantification images from Figure 6A-6B were collected using the Zeiss LSM800 confocal with an EC Plan-Neo Fluor 1.25x/0.03 and Plan-Apochromat 63x/1.4 lens, respectively.

#### Image analysis

All images were analyzed and prepared for display using FIJI ^68^. Brightness and contrast were adjusted on a per image basis in order to best display cell positions and identifications, unless otherwise noted (e.g. CellROX intensity in biofilms imaged and displayed the same for comparison of intensity (Figure 6A-6B)). For time lapse imaging, agar pad XY drift was accounted for by performing an image registration in FIJI using the StackReg plugin with the Rigid Body option. For CFU counting and aggregate biofilm size measurements, images were segmented following a background subtraction, contrast enhancement, and median filter. The processed fluorescence channels were segmented using the Auto Local Threshold function. Segmented objects were counted using the Analyze Particles function with a size filter set to the average size of the specified objects. The XY image position was recorded and used to find the Euclidian distance between aggregate biofilms on a per-image basis. Empirical Cumulative Distribution Function (ECDF) plots (Figure 3F and Figure 6G) display 95% confidence intervals and were generated in Python using iqplot ^69^.

CellROX intensity was measured from a subset of the images in Figure 6A-6B by generating an intensity-based threshold binary mask of the consumer space (GFP channel). The producer space was then found by inverting the consumer space. Finally, the producer remaining was found by thresholding the producer channel (mApple). By finding the difference between the producer space and producer remaining, a percentage was calculated (Figure 6D, Figure S4B). Finally, to assess the distance in which the producer was saved from or induced consumer lysis, the intensity profile of a line was drawn between the two points and used to mark start and end and recorded for several images.

### QUANTIFICATION AND STATISTICAL ANALYSIS

Quantification and analyses are described in the Method Details. Briefly, experimental values were compiled and explored in Python and results were plotted using either Seaborn, iqplot, or exported and displayed in GraphPad Prism v9. Statistical tests were performed in GraphPad Prism v9, and the relevant statistical test and sample size is described in the appropriate figure legend.

