## Supplementary material for "The contrasting roles of nitric oxide drive microbial community organization as a function of oxygen presence": Figures S1-S4, Supp Table 1

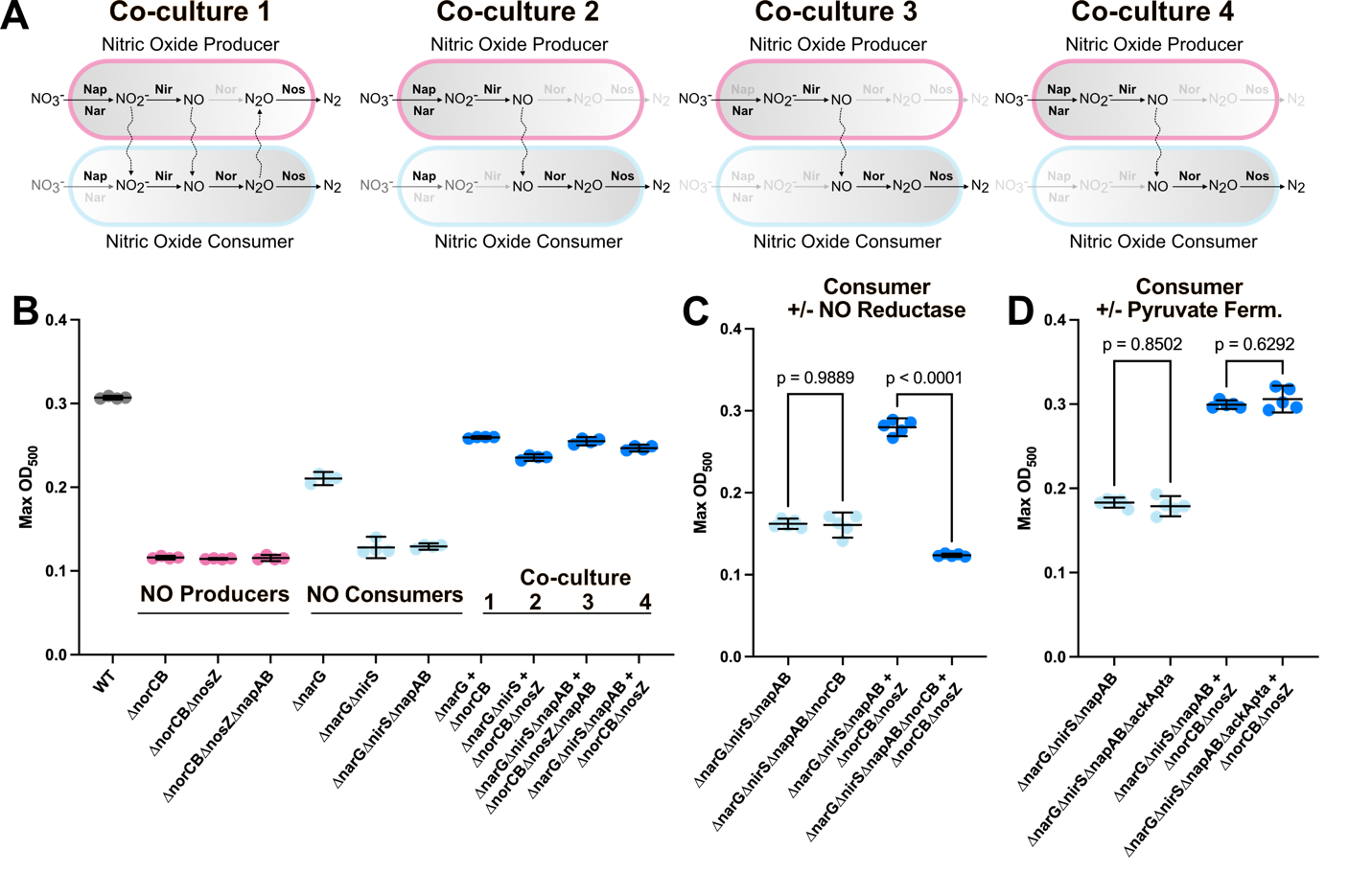

**Figure S1**. **Synthetic NO cross-feeding co-culture comparisons.** **Related to Figure 2.** (**A**) Schematic representation of genomic identity and subsequent cross-feeding of different NO producer and consumer strains. Wiggly arrows indicate intermediates that can be exchanged. Grayed-out gene name indicates a clean genomic deletion. (**B**) Max OD for specified strains/co-cultures from growth curves performed identically to those in Figure 2B. (**C**) Comparison of consumer and co-culture growth when consumer possess the nitric oxide reductase (Nor). **D**) Comparison of consumer and co-culture growth when consumer possess the ability to generate ATP via pyruvate fermentation (AckApta). (**B-D**) Max OD reached over the course of 72-hour growth curves supplemented with 5mM nitrate. Each dot is an independent well (technical replicate, n=4 (B) or n=5 (C,D) for each condition). Each graph is from an independent experiment. Line and error bars in graphs indicate mean and 95% confidence intervals; P values reflect one-way ANOVA with Tukey’s multiple comparisons test.

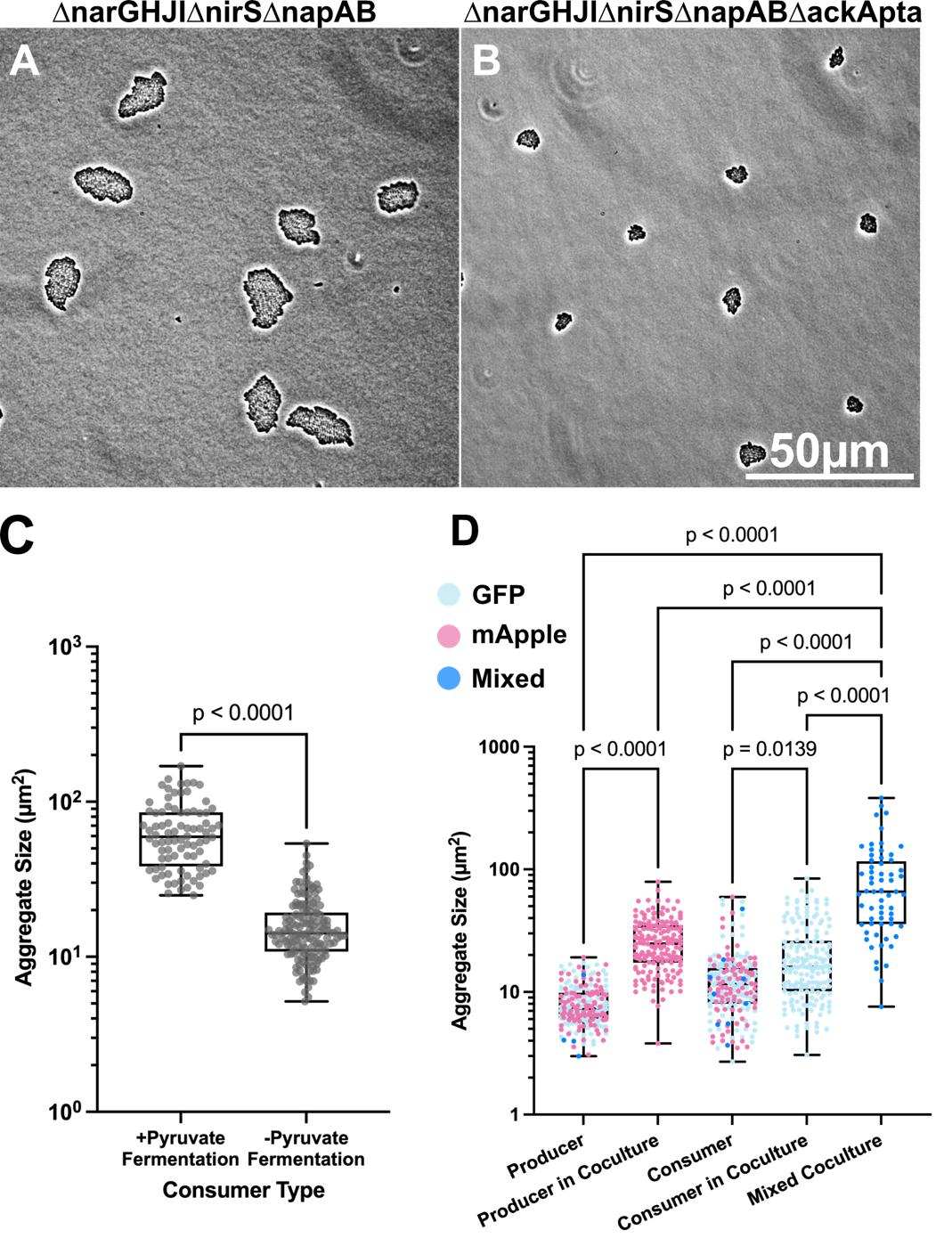

**Figure S2.** **Single cell aggregate imaging increases our ability to observe subtle slow-growth differences. Related to Figure 3.** (**A-B**) Comparison of consumer strains differing only in their ability to perform pyruvate fermentation (ackApta) grown under agar pads as in Figure 3B. (**C**) Quantification of aggregate sizes. P value from unpaired t-test. Each point represents one aggregate (n=84 consumer + pyruvate fermentation, n=153 consumer – pyruvate fermentation) measured from 10 images per condition, (**D**) Breakout of aggregate size quantification from Figure 3E. P values reflect one-way ANOVA with Tukey’s multiple comparisons test.

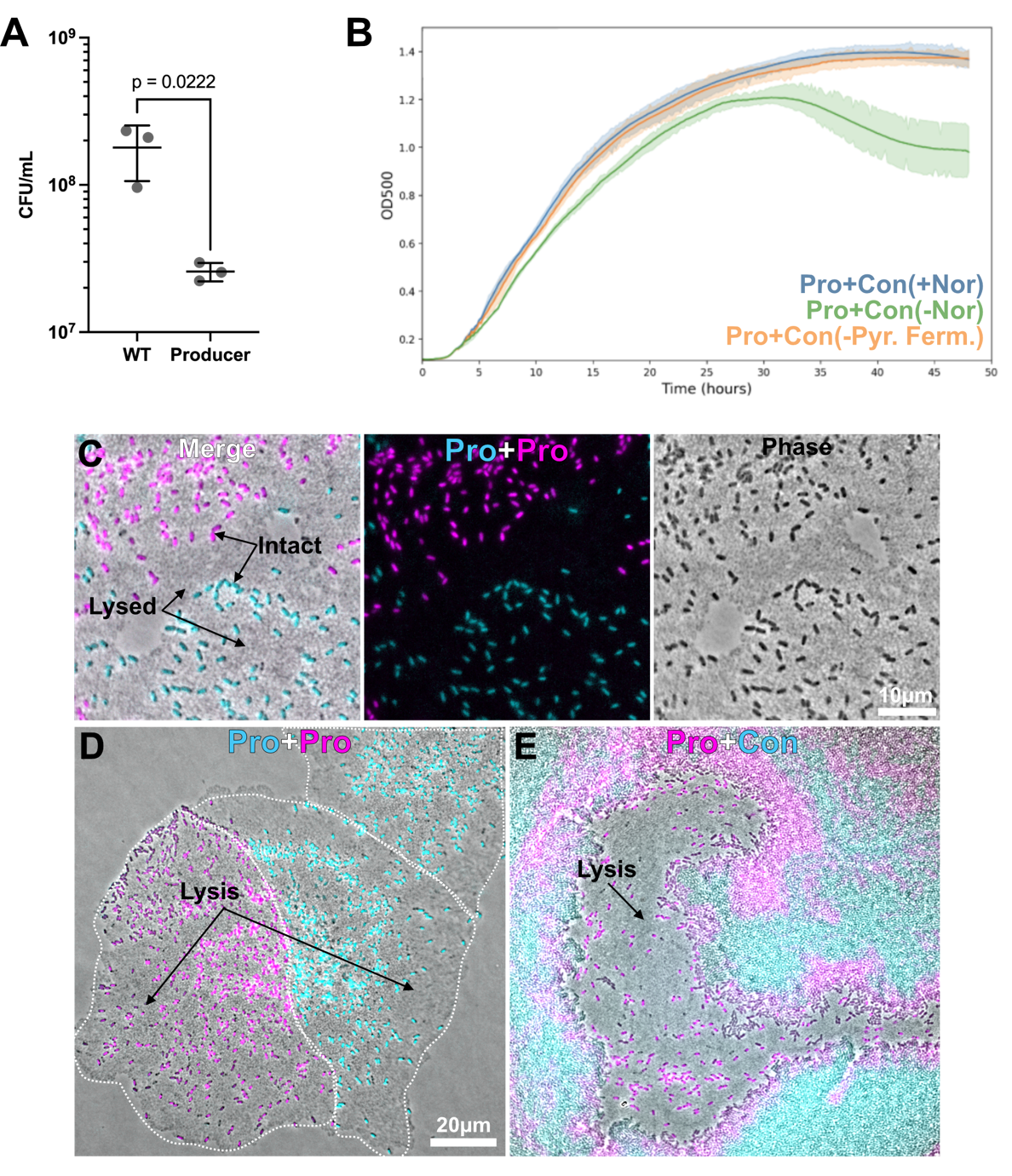

**Figure S3. Confirming the cause of reduction in OD under hypoxic growth conditions.** **Related to Figures 4 and 5.** (**A**) Specified strains grown under identical conditions to Figure 4F (hypoxic-not shaking, with nitrate) were plated for colony forming units (CFU). NO producer strain reduction in CFU is consistent with observed reduction in OD and cell death interpretation. Each point represents one replicate (n=3 per condition). Line and error bars indicate mean and standard deviation; P value from unpaired t-test. (**B**) Comparison of NO producer co-cultured with different consumers. A rescue from cell death (reduction in OD) during co-culture is not observed when consumer lacks NO reductase (Nor)). Each line represents the mean of n≥4 replicates per condition. (**C**) NO Producer comparison of fluorescence and phase channels used to interpret visual lysis. (**D-E**) Fluorescence and phase channel overlay of Pro+Pro and Pro+Con representative images in Figure 5.

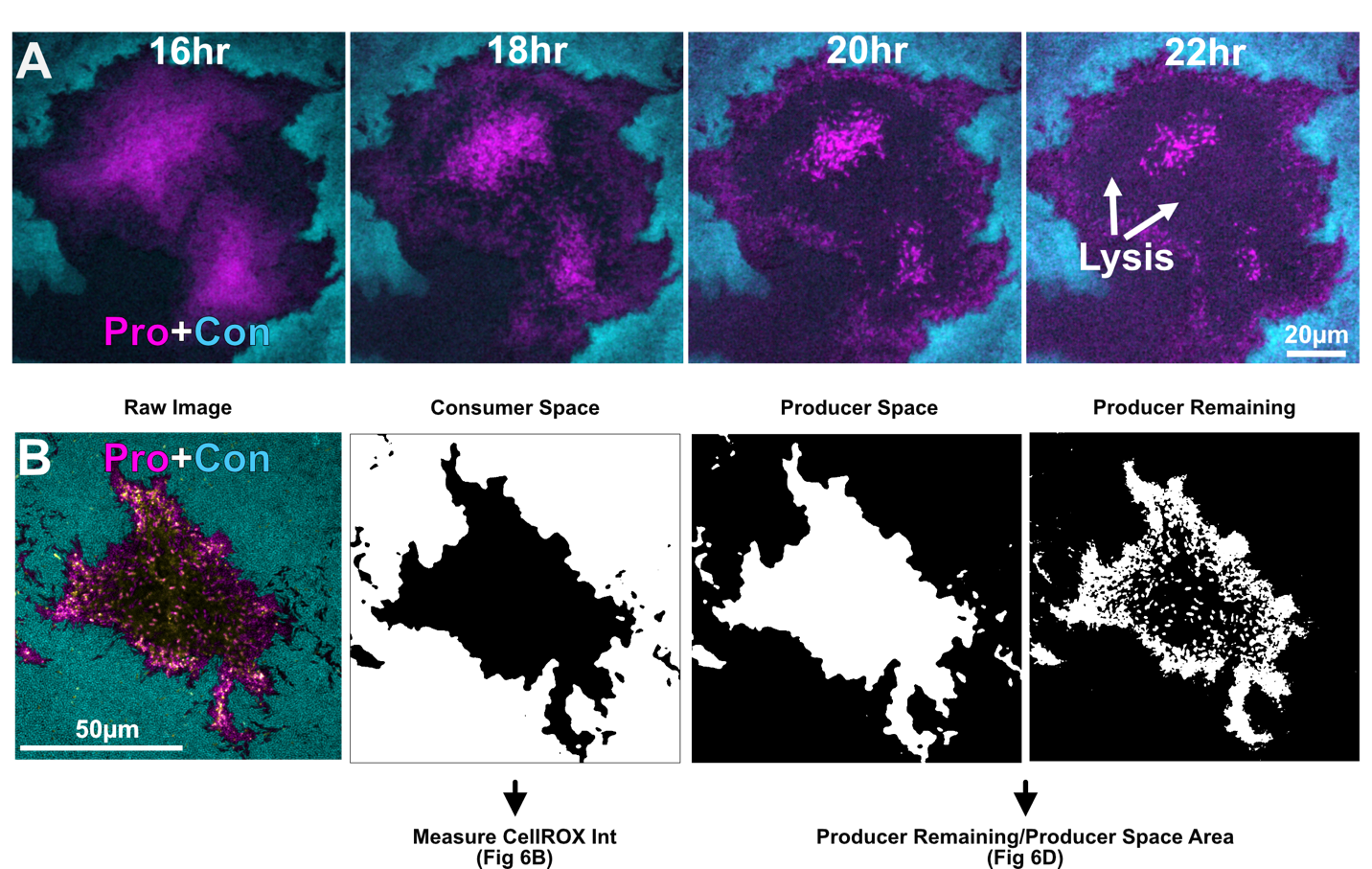

**Figure S4. Full patch time lapse and image analysis example. Related to Figures 5 and 6.** (**A**) Time lapse of a NO producer patch showing temporal lysis patterning occurring in the absence of consumer contact. (**B**) Example of image analysis masks used to quantify metrics in Figure 6 panels C and D.

| **Deletion Construct** | **NarGHJI** | **NapAB** | **NirS** | **NorCB** | **NosZ** | **AckA-Pta** |
| --- | --- | --- | --- | --- | --- | --- |
| **Function** | Membrane bound nitrate reductase | Periplasmic nitrate reductase | Nitrite reductase | Nitric oxide reductase | Nitrous oxide reductase | Pyruvate Fermentation |
| **Up F** | TAAAACGACGGCCAGTGCCACGTACTGGGTGTTCGCCCTG | ACGACGGCCAGTGCCAAGCTTCTACAGTCGCCTGCATAGA | ACGACGGCCAGTGCCAAGCTTGTCGATGCGCTGGGCCAGCA | TAAAACGACGGCCAGTGCCATGATCCTCGGCTGGCTGG | ACGACGGCCAGTGCCAAGCTACGTACTTGACCTGCTGGCG | ACGACGGCCAGTGCCAAGCTTTCAGGCTGCAGAAGGACTG |
| **Up R** | CGCGCAGGGTCTTGATCTCCTCACCCGGTC | GTGCTTTCATGCGGCCTCCCCGATTGTCTCCTCGGGCGAG | GGCTCCTGAGGAGATAGACCGACCCGCGTGCGGGGCACGC | GAAGGCCATCGCGGCCTCCTGGAATGGG | GCCCTTGAGGACGACACGAGTCCACTCAGCGCGGTCGATG | GCCCACTGGGCGGCGTTCCTTCACTGCTCCTTGGTCTGCT |
| **Dn F** | GGAGATCAAGACCCTGCGCGGCCGGCGCAT | CTCGCCCGAGGAGACAATCGGGGAGGCCGCATGAAAGCACTCA | GCGTGCCCCGCACGCGGGTCGGTCTATCTCCTCAGGAGCC | AGGAGGCCGCGATGGCCTTCGCCGTCGAG | CATCGACCGCGCTGAGTGGACTCGTGTCGTCCTCAAGGGC | AGCAGACCAAGGAGCAGTGAAGGAACGCCGCCCAGTGGGC |
| **Dn R** | CATGATTACGAATTCGA GCTGCTGGCGCGGCAGGAAGCGC | CATGATTACGAATTCGAGCTTCCTGCCGACCGGGGCCCAG | CATGATTACGAATTCGAGCTTATCAGGCGACATGGAGATC | CATGATTACGAATTCGAGCTCCGGGCAATCGTCCAGCG | CATGATTACGAATTCGAGCTGAGGAACTGGCCGCCCTGGA | CATGATTACGAATTCGAGCTTACTGATCGCGGCCTGGAAGAAAAAGC |
| **Plasmid** | pMQ30 | pMQ30 | pMQ30 | pMQ30 | pMQ30 | pMQ30 |
| **Cut Sites** | HindIII, SacI | HindIII, SacI | HindIII, SacI | HindIII, SacI | HindIII, SacI | HindIII, SacI |
| **Published** | Spero et al 2018 | This study | This study | This study | This study | This study |

| **Strain name** | **Parent** | **Deletion** | **Function of deletion** | **Published** |
| --- | --- | --- | --- | --- |
| **PA14 (WT)** | PA14 | NA | NA | NA |
| **∆narGHJI** | PA14 | NarGHJI | Membrane nitrate reductase | Spero et al 2018 |
| **∆norCB** | PA14 | NorCB | Nitric oxide reductase | This study |
| **∆nirS** | PA14 | NirS | Nitrite reductase | This study |
| **∆narGHJI∆nirS** | ∆nirS | NarGHJI | Membrane bound nitrate reductase, nitrite reductase | This study |
| **∆norCB∆nosZ** | ∆norCB | NosZ | Nitric oxide reductase, nitrous oxide reductase | This study |
| **∆narGHJI∆nirS∆napAB** | ∆narGHJI∆nirS | NapAB | Membrane nitrate reductase, nitrite reductase, periplasmic nitrate reductase | This study |
| **∆norCB∆nosZ∆napAB** | ∆norCB∆nosZ | NapAB | Nitric oxide reductase, nitrous oxide reductase, periplasmic nitrate reductase | This study |
| **∆narGHJI∆nirS∆napAB∆norCB** | ∆narGHJI∆nirS∆napAB | NorCB | ∆narGHJI∆nirS∆napAB - NO reductase ability | This study |
| **∆narGHJI∆nirS∆napAB∆ackApta** | ∆narGHJI∆nirS∆napAB | ackApta | ∆narGHJI∆nirS∆napAB - Substrate level phosphorylation via pyruvate fermentation ability | This study |

**Supplementary Table 1**. **Primers, plasmids, cut sites and strains used to make the**

**mutants used in this study. Related to Figure 2.**
